# The impact factor fallacy

**DOI:** 10.1101/108027

**Authors:** Frieder Michel Paulus, Nicole Cruz, Sören Krach

## Abstract

The use of the journal impact factor (JIF) as a measure for the quality of individual manuscripts and the merits of scientists has faced significant criticism in recent years. We add to the current criticism in arguing that such an application of the JIF in policy and decision making in academia is based on false beliefs and unwarranted inferences. To approach the problem, we use principles of deductive and inductive reasoning to illustrate the fallacies that are inherent to using journal based metrics for evaluating the work of scientists. In doing so, we elaborate that if we judge scientific quality based on the JIF or other journal based metrics we are either guided by invalid or weak arguments or in fact consider our uncertainty about the quality of the work and not the quality itself.

## Introduction

The journal impact factor (JIF) was initially used to help librarians make decisions about journals (Garfield, 2006). However, during the last decades the usage of the JIF has significantly changed. In deviating from its original purpose it is now widely used to evaluate the quality of individual publications and the work of scientists (Amin & Mabe, 2003; Arnold & Fowler, 2010). Since then, the measure itself has been criticized for various reasons. For example, it is well known that the JIF is an inaccurate estimate for the expected number of citations of an article within a specific journal (Callaway, 2016; Lariviere et al., 2016) and that it is relatively easy to manipulate (McVeigh & Mann, 2009; Tort, Targino, & Amaral, 2012). Nonetheless, the JIF has deeply affected the work of scientists and decision making in academia. Scientists get jobs, tenure, grants, and bonuses based on the impact of the journals they are publishing their manuscripts in, outgrowths’ which were critically discussed in many previous reviews, comments and editorials (Casadevall & Fang, 2014; Della Sala & Crawford, 2007; DePellegrin & Johnston, 2015; Lehmann, Jackson, & Lautrup, 2006; Reich, 2013; Seglen, 1997; Simons, 2008; Werner, 2015, and please see Brembs, Button, & Munafò, 2013 for very thorough analyses of the detrimental effects of the JIF). Notably, the JIF has also been explicitly referred to as a tool to decide how to distribute funds across institutions, for example in Germany (German Science Foundation [DFG], 2004), and thereby affects policy making on a much larger scale.

> "For the calculation of the performance-based bonus of the unit providing the service (department or clinic) the original publications may be used with the unweighted impact factor of the publication organ, in the sense of a step-wise introduction of quality criteria. Thereby, a first- and last authorship may be considered with one third each and the remaining third can be distributed across all remaining authors […]."^1^

Besides such explicit usage of the JIF for evaluating scientific excellence, the JIF also implicitly affects other measures which have been suggested to better approximate the quality of a scientist's work or of a specific study (e.g. the h-index, Hirsch, 2005 and the Relative Citation Ratio (RCR), Hutchins, Yuan, Anderson, & Santangelo, 2015). For example, there is some evidence that the number of citations of an article is influenced by the JIF of the journal where the article was published, regardless of the quality of the article itself (Brembs et al., 2013; Callaham, Wears, & Weber, 2002; Cantrill, 2016; Lozano, Larivière, & Gingras, 2012). This implies that measures that are based on the citations of the individual articles are still influenced by the JIF of the publication organ. With the many different ways of how the JIF can influence decision making in academia, it is not surprising that empirical data now demonstrate the JIF to be one of the most powerful predictors for academic success (Van Dijk, Manor, & Carey, 2014).We could recently show that some scientists may have adapted to these reward principles in their environment by showing a greater reward signal in the brain’s reward structures in the prospect of an own high impact publication (Paulus, Rademacher, Schäfer, Müller-Pinzler, & Krach, 2015).

In line with the rising initiatives to prevent the use of the JIF for evaluating the quality of science (see e.g. the DORA initiative, Alberts, 2013, Cagan, 2013 or the report of the (German Council of Science and Humanities [Wissenschaftsrat], 2015), we have considerable doubts that the arguments in support of using the JIF for measuring scientific excellence are justified. In this comment we want to look at the problem of using the JIF from a different perspective and carefully (re)evaluate the arguments for its use as an estimate of scientific quality. Thereby, we hope to better understand the beliefs about the JIF that influence decisions in academia and the implications of policies that use the JIF to assess and remunerate scientific quality. Beyond the specific case of the JIF, this exercise might also help to specify more general misconceptions when using journal based properties to evaluate science, in order to overcome incentive structures based on journal based metrics altogether.

## Deductive fallacy when using the JIF

A basic belief when using the JIF for evaluating the quality of a specific manuscript seems to be that (1) if a paper is published in a high impact factor journal (p) then the paper is of high quality (q) ^2^. Why would scientists believe this? A straightforward reason is the idea that it is more difficult to publish in a high impact factor journal because higher standards of research quality and novelty have to be passed in order to be accepted. The average number of citations of a journal's articles within in a specific time period signals the average breadth of interest in these articles during that time period, which can of course be affected by many factors other than research quality. But as a first approximation, let us suppose that belief (1) is the case. What can we conclude from it? If we see a paper published in a high impact factor journal, we could then draw the deductively valid inference of modus ponens (MP: *if p then q, p, therefore q*)^3^ and conclude that the paper is of high quality. But what if we see a paper published in a low impact factor journal? Can we draw any conclusions in this case?

One aspect of the impact factor fallacy could be operationalized as the tendency to draw the deductively invalid inference of *denial of the antecedent* (DA: *if p then q, not-p, therefore not-q*). This inference is deductively invalid because it is logically consistent for the premises *if p then q* and *not-p* to be true and yet the conclusion *not-q* to be false. When the premises of an inference can be true and at the same time the conclusion false, the inference does not preserve truth when going from premises to conclusion. In order to argue that the conclusion is not false in a particular case, we would therefore have to go beyond this argument and provide further information that might increase support for the conclusion.

For the more realistic case that the premises and conclusion are uncertain, such that they can not only be either true or false, but can be held with varying degrees of belief, the inference of DA is probabilistically invalid (p-invalid) because there are coherent^4^ probability assignments to premises and conclusion for which the probability of the conclusion is lower than the sum of the probabilities of the premises (Adams, 1998; Over, 2016). Therefore, just like in the binary case DA does not preserve truth from premises to conclusion, in the probabilistic case DA does not preserve probability from premises to conclusion, so that it would be warranted to have a high degree of belief in the premises and yet a very low degree of belief in the conclusion. In order to justify the conclusion in a particular instantiation of the argument, we would have to bring further information into the discussion beyond that contained in the premises. Applied to the JIF example, suppose we assume that if a paper is published in a high impact factor journal, it is of high quality, and then encounter a paper that is published in a low impact factor journal. From this alone it is not justified to conclude that the paper we encountered is not of high quality. In order to draw such a conclusion we would require more information.

Denial of the antecedent (DA) is of course not the only inference one can draw on the basis of the conditional belief that if a paper is published in a high impact factor journal, then it is of high quality. A similar, deductively valid inference results if we add a further premise to DA: "If a paper is not published in a high impact factor journal, then it is not of high quality". One can combine this new conditional premise with the conditional premise that we already had: "If a paper is published in a high impact factor journal, then it is of high quality", to obtain the following biconditional premise: "A paper is published in a high impact factor journal if and only if it is of high quality". From this biconditional premise (or equivalently from the two conditional premises) together with the premise that a specific paper was not published in a high impact factor journal, one can indeed validly conclude that the paper is not of high quality. However, this inference will only be useful if one believes the biconditional premise to a non-negligible degree in the first place. If the biconditional premise is implausible, then any deductively valid conclusion based on it will also tend to be implausible, precisely because it follows logically from an implausible starting assumption. Considering that most scientists are likely to agree that it is not only implausible but false that a paper is of high quality if and only if it is published in a high impact factor journal, the fact that the inference from this biconditional is valid has no use for practical purposes.

## Inductive fallacies when using the JIF

One could argue that deduction, and with it logical validity, has little impact on actual reasoning and decision making outside of the mathematics classroom, and that therefore the inferences we should be looking at when analysing the use of the JIF in the practice of science should rather be inductive (Baratgin & Politzer, 2016; Chater, Oaksford, Hahn, & Heit, 2011; Evans, 2002; Oaksford & Hahn, 2007).

An inductive inference that might describe well the use of the impact factor is the informal fallacy of the *argument from ignorance* (or its Latin equivalent "ad ignorantiam"). This argument tries to justify a conclusion by pointing out that there is no evidence against it. Typical examples could be "No side effects were found for this treatment in clinical trials. Therefore this treatment is safe" or "No one has proven that ghosts do not exist. Therefore ghosts exist" (Hahn & Oaksford, 2007; Oaksford & Hahn, 2004, 2007). In the case of the JIF, if a paper comes from a high impact journal this can be seen as a sign suggesting it is an excellent piece of work. But as we saw above in the discussion of DA, this does not imply that if the paper was published in a low impact factor journal this is a sign suggesting that the quality of the paper is low. A more precise description of the situation would be that a low impact factor journal lacks the sign of high quality that a high JIF provides. If a paper is published in a low impact journal then we have less information about its quality, rather than having information suggesting that its quality is low. It is an argument from ignorance to use the absence of impact factor based evidence for high quality to conclude that a paper is of low quality.

However, the argument from ignorance is not always a bad argument (Hahn & Oaksford, 2007, 2012). Its strength depends on how informative the lack of information about something being the case is in the situation at hand. Suppose we search a book in a library catalogue and do not find it. In this case it is reasonable to use the lack of information about the book in the catalogue to conclude that the book is not in the library. Similarly, if we look at a train timetable and do not see a particular town listed, it is reasonable to conclude that the train does not stop in that town. However, suppose we are planning a party and have invited the whole department, in the hope that a particular person we are attracted to will attend. In this case a lack of information indicating that the person will come does not warrant the conclusion that the person will not come. Catalogues and timetables are fairly closed environments in which we can expect all relevant information to be stated explicitly. But environments like those of social interactions or research endeavours are typically more open, so that the absence of information about something being the case simply does not warrant us to conclude that it is not the case. A consequence for the JIF would be that low impact publications do not signal low research quality, but rather uncertainty about the quality and the need to gather more information in order to be able to determine research quality.

Two further inductive inferences that might be relevant in accounting for the use of the JIF are the informal fallacies of the *argument from authority* (also called by the Latin name "ad verecundiam"), and of the *ad hominem argument* (Bhatia & Oaksford, 2015; Hahn & Hornikx, 2016). The argument from authority tries to justify a conclusion by pointing out that some expert or authority endorses the conclusion. Typical examples could be "Scientist x says that the treatment is safe. Therefore the treatment is safe", "My parents say that Santa Claus exists. Therefore Santa Claus exists" or "My peers say that clothing item x is great. Therefore clothing item x is great". In the case of the JIF, a high impact factor of a journal would play the role of an authority for the quality of the papers within it.

In contrast, the ad hominem argument tries to justify the rejection of a conclusion by pointing to personal attributes of a person that endorses it. Typical examples could be "The new treatment was developed by a person with no formal degree in the subject. Therefore the treatment is not safe", or "A person without a driver’s license says “don’t drink alcohol while driving". Therefore, it is false that you should not drink alcohol while driving”. In the case of the JIF, a low impact factor would be used to give a journal a reputation of low quality, and this low quality reputation would then be transferred to the papers within it.

Like the argument from ignorance, the argument from expert opinion and the ad hominem argument are not always bad arguments. Their quality varies as a function of how informative the authority status, or the personal attributes of the instance endorsing them, is for the problem at hand. Policy decisions are routinely based on the advice of experts, and there seems to be agreement that this is a good thing to do, as long as the experts are really considered experts in their field and their advice is not biased (Harris, Hahn, Madsen, & Hsu, 2016; c.f. Sloman & Fernbach, 2017). Dismissing an argument because of personal attributes of a person endorsing it is often more difficult, because it has to be made plausible that those attributes are relevant to the quality of the argument. For example, that one does not need to be a mother to be qualified for being prime minister seems obvious, whereas a case of a person applying to a position against gender discrimination, who in his private life beats his wife, is likely to be more controversial. In the case of the JIF, we would have to justify why we think that a low impact factor indicates that a particular journal is of low quality, and why this low quality can be transferred to a particular paper within it. Such a judgment requires further information about the journal and about the paper at hand to be justified, which is usually not provided. Thus, whereas a high impact factor may add to the reputation of a journal, a low impact factor does not warrant a bad reputation, but rather provides insufficient information about reputation (see Table 1 for examples of the inductive and deductive fallacies as discussed here).

**Table 1.** The deductive and inductive fallacies discussed in this paper.

| Name | Form | Plausible Example | Implausible Example | Journal Impact Factor Example |
| --- | --- | --- | --- | --- |
| **Deductive fallacy** |  |  |  |  |
| Denial of the antecedent | *If $p$ then $q$. Not-$p$. Therefore not-$q$.* | If the glass falls down then it breaks. The glass does not fall down. Therefore, the glass does not break. | If you carry an umbrella then you stay dry. You do not carry an umbrella. Therefore, you do not stay dry. | If a paper is published in a high impact factor journal, then it is of high quality. This paper is not published in a high impact factor journal. Therefore, this paper is not of high quality. |
| **Inductive fallacies** |  |  |  |  |
| Argument from ignorance | *It is not known that $p$ is true (false). Therefore $p$ is false (true).* | The book is not listed in the library catalogue. Therefore, the book is not in the library. | No one has proven that ghosts do not exist. Therefore, ghosts exist. | This paper does not have the quality sign of having been published in a high impact factor journal. Therefore, this paper is not of high quality. |
| Argument from authority | *This expert says that $p$ is true. Therefore $p$ is true.* | Medical experts say that this treatment is safe. Therefore, this treatment is safe. | My parents say that Santa Claus exists. Therefore, Santa Claus exists. | This paper does not have the authority backing of having been published in a high impact factor journal. Therefore, this paper is not of high quality. |
| Ad hominem argument | *This untrustworthy person says that $p$ is true. Therefore $p$ is false.* | A person without training says that this treatment is safe. Therefore, this treatment is not safe. | A person without a driver's license says "don't drink alcohol while driving". Therefore, it is false that you should not drink alcohol while driving. | This paper was published in a journal with low quality reputation due to a low impact factor. Therefore, this paper is not of high quality. |

Until now we have discussed inferences on the basis of the belief that if a paper is published in a high impact factor journal, then it is of high quality. But although this belief can sometimes be useful as a quick approximation or rule of thumb, it is often itself not warranted. Not only because the mean number of citations of the papers in a journal is an indicator of the average breadth of interest in these papers during the first years after publication, which is not the same as research quality (e. g. a high quality paper may have low citation rates because it is addressed to a small, highly specialised audience, or because its significance is only realised five years after publication; and a paper may have high citation rates because of highly consequential flaws within it). But more specifically, it is often not warranted because the inference from a metric defined at the journal level to the features of an individual paper within that journal involves an ecological fallacy.

## Ecological fallacy when using the JIF

Finally, the evaluation of manuscripts based on the JIF bears an ecological fallacy. When comparing group level data, such as the average citations of journals, it is difficult up to impossible to infer the likelihood of the outcome for comparisons on the individual level, such as citations of manuscripts. In fact, it is relatively easy to think of examples where the likelihood to find a manuscript with more than twelve citations per year in a lower impact journal exceeds the likelihood of finding such manuscript in a higher impact journal. This type of ecological fallacy occurs when the distribution of citations is heavily and differentially skewed within each higher level unit, i.e. the journals. This is typically the case when it comes to citation rates of journals (see e.g. Lariviere et al., 2016). Accordingly, a journal with a JIF of twelve might contain few manuscripts that were cited several hundred times in the previous two years, but many others that were not cited at all during the same period. Such a citation pattern would result in a heavily skewed distribution of citations per article, while another journal with a JIF of ten might have a normally distributed citation rate of articles for the same time period. Without further knowledge of the distribution of citations within the journals in a given year (i.e. information at the individual level) concluding that a manuscript in the journal with a higher JIF is of better quality (or of broader interest) involves an ecological fallacy, because it is possible that the likelihood of finding a manuscript with more citations in the lower impact journal is in fact similar or even higher.

## Concluding remarks

With this comment, we hope to have highlighted some misconceptions in the beliefs and arguments involved in using journal based metrics, and specifically the JIF, for evaluating the work of scientists. While some of the thoughts described here are introduced to illustrate the most controversial arguments, others better approximate the reality of decision making in academia. In this exercise, it is surprising to see many political and academic institutions as well as scientists having believed for so long that they are evaluating the "quality of science" while they are keen to provide weak arguments, draw invalid conclusions, or weigh their lack of information and uncertainty about the subject when using the JIF.

From an economic perspective, however, it might in fact be a successful strategy to minimize the uncertainty about the quality of the evaluated work, person, or institution by relying on the JIF, and it might also be better to have a weak argument than to have no argument. Evaluating the quality of a scientist’s work surely is a time-consuming process and it takes much more effort than simply comparing impact factors. Accordingly, deans, commissions, or institutions which might not have the resources for an actual assessment of "scientific excellence" have reasons to rely on the JIF. However, it should be clear that those decisions are not based on the *quality* of the scientific contribution per se but, optimistically, somehow integrate the *availability of information* about the quality. This distinction makes an important difference for communicating and justifying decisions in academia. As an illustrative example, one can compare the situation of deciding that a candidate does not deserve tenure because one thinks that the quality of the work was not good enough, to deciding that a candidate does not deserve tenure because one lacks information and is uncertain whether the quality of the work was good enough. While persons and institutions usually *communicate* as if they were following the first argument, their *justification* most often implies the latter if they base their decisions on journal based metrics.

The JIF is arguably the most popular journal based metric of our times, but it has already been subject to severe criticism in the past (Brembs et al., 2013; Della Sala & Crawford, 2007; DePellegrin & Johnston, 2015; Lehmann et al., 2006; Reich, 2013; Seglen, 1997; Simons, 2008; Werner, 2015). As a result, it seems that (some) individuals and institutions within the scientific community are ready to shake off the JIF at some point in the nearer future (Alberts, 2013; Cagan, 2013; Callaway, 2016). We want to point out that the problems described here apply in one way or another to any journal based assessment. If journals would drop out of the ‘impact factor game’ (PLoS Medicine Editorial, 2006) publications in some journals might still be regarded as more valuable than in others. It is difficult to quantify those influences, but having a publication in one of the ‘golden club’ journals (Reich, 2013) could simply replace the metric of the JIF with another, more implicit qualitative measure for distinguishing prestigious from less prestigious journals. Thereby, the fallacies and problems described above would continue to govern decision making in academia as long as we base them on any kind of journal based assessment and the rank of publication organs.

## Author contributions

FMP, NC, and SK wrote the manuscript.

## Acknowledgments

We would like to thank Inês Mares and Mike Oaksford for helpful comments and discussion on an earlier draft of this manuscript.

## Footnotes

1 “Für die Berechnung der LOM [leistungsorientierte Mittel; remark of authors] der jeweiligen leistungserbringenden Einheit (Abteilung bzw. Klinik) kann im Sinne einer stufenweisen Einführung von Qualitätskriterien die Bewertung erfolgter Original-Publikationen unter Verwendung des ungewichteten Impact Faktor der jeweiligen Publikationsorgane (JIF) erfolgen. Dabei können Erst- und Letztautorschaft mit je einem Drittel berücksichtigt werden; das verbleibende Drittel kann auf alle übrigen Autoren verteilt werden […]." (German Science Foundation [DFG], 2004, p. 15).

2 When we speak of “high” and “low” impact in this paper, the arguments we make are independent of whether “high” and “low” refer to the absolute JIF of a journal, or to the JIF relative to a specific research domain.

3 Here *p* and *q* stand for arbitrary propositions. For example, *p* might stand for “This paper is published in a high impact factor journal” and *q* for “This paper is of high quality”.

4 Two statements are *coherent* if and only if they respect the axioms of probability theory. For example, these axioms state that if we believe it is 80% likely to rain, then in order for our beliefs to be coherent we should also be willing to believe that it is 20% likely not to rain, otherwise the probabilities involved would not sum up to 1.

